# Sleep-related consolidation and generalizability of motor skills learned by physical practice, motor imagery and action observation

**DOI:** 10.1101/2022.12.06.519290

**Authors:** Adrien Conessa, Ursula Debarnot, Isabelle Siegler, Arnaud Boutin

## Abstract

Sleep benefits the consolidation of motor skills learned by physical practice, mainly through periodic thalamocortical sleep spindle activity. However, motor skills can be learned without overt movement through motor imagery or action observation. Here, we investigated whether sleep spindle activity also supports the consolidation of non-physically learned movements. Forty-five electroencephalographic sleep recordings were collected during a daytime nap after motor sequence learning by physical practice, motor imagery or action observation. Our findings reveal that a temporal cluster-based organization of sleep spindles underlies motor memory consolidation in all groups, albeit with distinct behavioral outcomes. A daytime nap offers an early sleep window promoting the retention of motor skills learned by physical practice and motor imagery, and its generalizability towards the inter-manual transfer of skill after action observation. Findings may further have practical impacts with the development of non-physical rehabilitation interventions for patients having to remaster skills following peripherical or brain injury.

## Introduction

Repeated practice is critical for the learning and mastering of motor skills. Training procedures encouraging the ability to exploit the features of a learned skill for its transfer from one situation to another is fundamental across diverse contexts, such as in sports sciences and rehabilitation^1^. Although the common way to learn a movement is by performing the task physically (see ^2,3^ for reviews), other forms of practice can contribute to motor-skill learning. Compelling evidence shows that motor skills can be learned without overt movement through motor imagery (MI) or action observation (AO)^4–7^. While MI refers to the process of mentally rehearsing a movement without physically performing it, AO consists in observing another actor performing the movement. Numerous neuroimaging studies reported that neural structures are commonly activated during MI, AO, and physical practice (PP), thus providing evidence of a relative “functional equivalence” between practice modalities^8–10^.

Both covert modalities of practice (MI, AO) engage a cognitive demand upon sensorimotor networks, boosting activity-dependent neuroplasticity and enhancing motor performance and learning^4,5,11^. Traditionally, to evaluate the beneficial effects of MI and AO practice on motor skill learning, participants observe a model or imagine themselves performing the motor task before being evaluated on a post-training test requiring the overt practice of the movement^5,12^. It is commonly reported that MI and AO practice lead to enhanced motor performance and learning, albeit to a lesser degree than PP^13–16^. However, motor learning is not limited to task-specific learning but also concerns the ability to transfer or generalize the newly acquired skill to another one or another effector (i.e., inter-limb transfer), which may depend on the modality of practice^1^. For instance, while the amount of PP influences the development of an effector-specific or unspecific representation of the sequence^17^, PP has been shown to rely more on an effector-specific representation of the motor skill after extensive practice than the two other modalities^5,18^, thus leading to impaired inter-limb skill transfer^17^. In contrast, MI and AO practice have mostly been revealed to develop an effector-unspecific representation of the learned motor task, thus allowing effective skill transfer from one limb to another^5,18^. Therefore, the representation of a motor skill acquired through PP, MI, and AO practice partly relies on distinct coding systems for movement production, leading to specific skill learning and transfer capacities^19–21^.

Although the repeated practice of a motor skill is crucial for its initial acquisition, the development of an effective movement representation is not only a result of practice^2^. The newly formed memory continues to be processed “offline” over several temporal scales, from a rapid form of consolidation at a microscale of seconds^22^ to longer forms of consolidation occurring primarily during the waking and sleeping hours following practice^2,23^. This offline period offers a privileged time window for memory consolidation, which relates to the process whereby newly acquired and relatively labile memories are transformed into more stable and enhanced memory traces^24^. The memory trace is thought to be dynamically maintained during wakefulness and actively reprocessed during a subsequent sleep period. A night of sleep and a daytime nap have been shown to play a crucial role in the strengthening and transformation of motor memory representations developed through PP during consolidation (see ^25^ for a review), which behaviorally results in either performance stabilization or improvement^26^. In the same way, a few studies reported sleep-dependent consolidation of motor skills learned through MI practice^11,12,27,28^ or AO^29^, and it has been recently emphasized that the stability of the newly formed motor memory through consolidation processes during wake and sleep episodes may modify its generalizability^1^. However, whether similar consolidation mechanisms are engaged during sleep following PP, MI and AO practice, and how sleep affects the generalizability of motor skills remains to be determined.

At the brain level, and in the context of PP, motor memory consolidation is thought to be mediated by transient thalamocortical sleep spindle activity – an electrophysiological hallmark of non-rapid eye movement stage 2 (NREM2) sleep involving brief 0.3-2 s bursts of waxing and waning 11-16 Hz oscillations. Sleep spindles have been suggested to support the offline reactivation of newly acquired motor memories, resulting in post-night and post-nap motor memory improvements^30–34^. Boutin and Doyon (2020)^25^ recently proposed a theoretical framework for motor memory consolidation following PP that outlines the essential contribution of the clustering and hierarchical rhythmicity of spindle activity during this sleep-dependent process. They posited that the rhythmic expression of sleep spindles over task-relevant cortical and subcortical brain regions is critical for the efficient reprocessing and consolidation of motor memory traces following PP. Specifically, it is suggested that the occurrence of sleep spindles follows two periodic rhythms: an infra-slow rhythm that corresponds to a 0.02 Hz periodicity of spindle-enriched periods, called spindle trains, and in which spindles iterate at an intermediate rhythm of about 0.2-0.3 Hz. Current theoretical models posit that this 0.2-0.3 Hz rhythmic occurrence of spindles during trains defines the sequential alternance of spindles and associated refractory periods, thus regulating the cyclic reinstatement and interference-free reprocessing of memory traces for their efficient consolidation^25,35–38^. However, whether a similar temporal cluster-based organization of sleep spindles underlies the consolidation of motor skills acquired by MI and AO practice remains to be addressed.

Hence, by combining behavioral and electroencephalographic (EEG) sleep measures, the aim of the present study was twofold: (i) to determine whether similar consolidation mechanisms are engaged during sleep following PP, MI and AO practice, and (ii) to investigate the specific contribution of sleep spindle activity and its rhythmic organization in motor skill consolidation and inter-manual transfer, depending on the modality of practice.

## Results

### Methods

#### overall experimental approach

Forty-five participants were required to learn a motor sequence task either by physical practice, motor imagery or action observation. Response times (i.e., interval between two keypresses; RT) were measured to evaluate skill performance during test blocks (Fig. 1): the pre-test and post-test blocks were respectively performed before and after the training phase, while the retention and inter-manual transfer tests were delayed and performed after a 90-min daytime nap following training. Motor skill acquisition is reflected by changes in RT performance (in percentages) from the pre-test to the post-test. Motor skill consolidation was assessed by analyzing the percentage of RT changes from the post-test to the retention test, while motor skill transfer was measured as the difference in RT performance (in percentages) between the retention test and the transfer test.

**Fig. 1.**
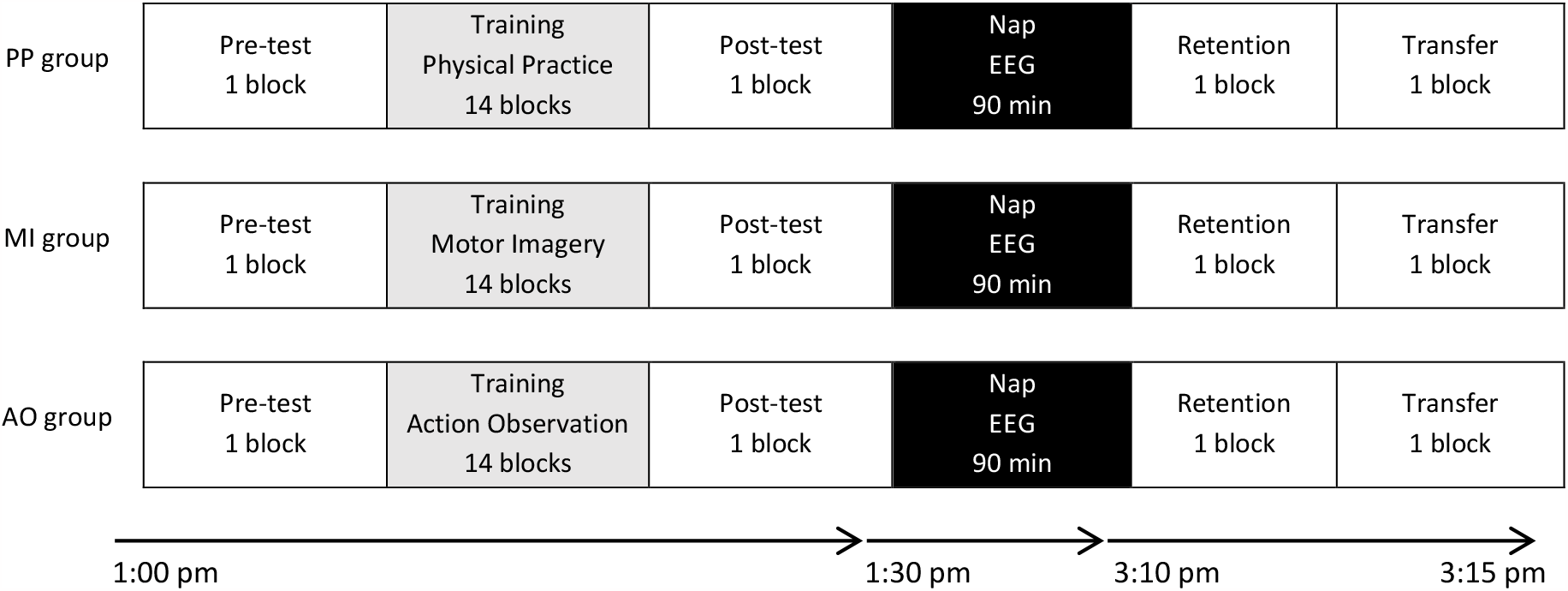
Experimental design. Participants (n = 45) were trained on a 5-element finger movement sequence either by physical practice (PP), motor imagery (MI) or action observation (AO). Sleep EEG recordings were acquired during a 90-min daytime nap following training. Participants were subsequently tested on the same motor sequence during retention and inter-manual transfer tests. During the pre-test, post-test, retention and transfer tests, all participants were required to physically performed the motor sequence. EEG: Electroencephalography.

#### Behavioral data

Fig. 2 illustrates the changes in performance associated with motor skill acquisition, consolidation and transfer capacity. A one-way ANOVA with the between-participant factor MODALITY (PP, MI, AO) was first performed on the RT data during the pre-test to ensure that the three groups did not differ at baseline performance. The analysis failed to detect a significant MODALITY effect (F(2, 42) = 2 .05, *p* = 0.14), revealing no performance differences between groups on the pre-test. Additional analyses on the RT data can be found in the supplementary material (see Fig. S1.). Then, one-way ANOVAs with the between-participant factor MODALITY (PP, MI, AO) were applied separately for motor skill acquisition, consolidation and transfer. For motor skill acquisition, the analysis revealed a main effect of MODALITY (F(2, 42) = 7.52, *p* = 0.002, *ɳ*^2^_p_ *=* 0.26), with better performance improvements for the PP group (M = 32.3%) in comparison to the MI group (M = 17.0%, *p* = 0.001). No significant difference in the magnitude of performance improvements was found between the AO group (M = 23.3%) and the other two groups (*p* = 0.06 and *p =* 0.12, respectively for the PP and MI groups). For motor skill consolidation, the analysis failed to detect a significant MODALITY effect (F(2, 42) = 2.50, *p* = 0.09). No significant difference in post-nap performance changes was found between the PP group (M = -0.2%), the MI group (M = 7.9%) and the AO group (M = 4.0%). For motor skill transfer, the analysis revealed a significant main effect of MODALITY (F(2, 42) = 5.10, *p* = 0.01, *ɳ*^2^_p_ *=* 0.20), with greater performance decreases for the PP group (M = -36.1%) than the MI group (M = -17.1%, *p* = 0.03) and the AO group (M = - 13.4%, *p* = 0.01), which did not significantly differ from each other (*p* = 0.63).

**Fig. 2.**
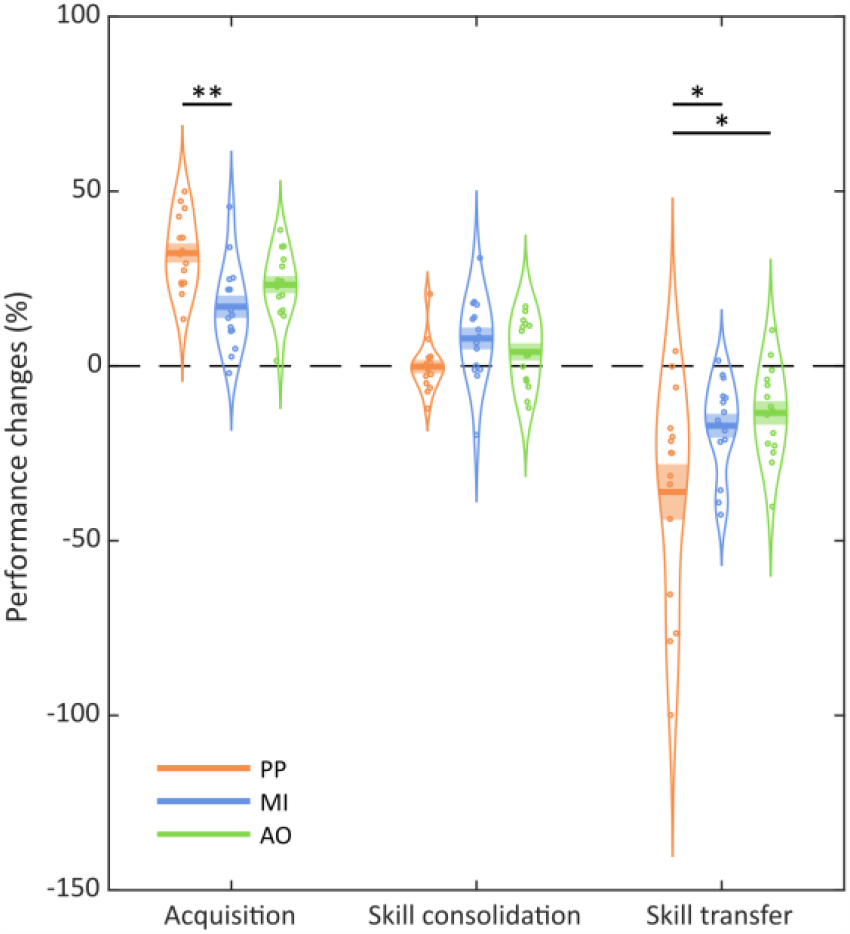
Behavioral results. Performance changes (in percentages) on the motor sequence task during acquisition (from the pre-test to the post-test), sleep-related skill consolidation (from the post-test to the retention test), and inter-manual skill transfer (from the retention test to the transfer test) for each of the physical practice (PP; n = 15), motor imagery (MI; n = 15) and action observation (AO; n = 15) groups. The curved lines indicate the distribution of data, the dark bars represent the mean of the distribution, and the lighter area surrounding the mean represent the standard error of the means. Individual data points are displayed as colored circles. * *p* < 0.05, ** *p* < 0.01.

#### Sleep and spindle characteristics

For all groups, sleep architecture and spindle metrics are respectively summarized in Table 1 and Table 2. Additional analyses for other scalp derivations (F3, F4, Fz, C3, C4, Cz, P3, P4, Pz, O1, O2, Oz) can be found in the supplementary material (see Fig S2. and S3). One-way ANOVAs with the between-participant factor MODALITY (PP, MI, AO) were applied separately for each spindle metric. *Spindle clustering*. The analysis did not reveal any significant main effect of MODALITY for the total number of sleep spindles (F(2, 42) = 0.22, *p* = 0.80), the number of grouped spindles (F(2, 42) = 0.12, *p =* 0.89) and isolated spindles (F(2, 42) = 0.81, *p* = 0.45), as well as for the number of spindle trains (F(2, 42) = 0.24, *p* = 0.80). *Spindle rhythmicity*. No significant main effect of MODALITY was found for the inter-spindle interval (ISI) within trains (F(2, 42) = 0.03, *p* = 0.97) and inter-train interval (ITI) metrics (F(2, 42) = 0.69, *p* = 0.51). *Spindle density*. The analysis did not reveal any significant main effect of MODALITY for both the global density (F(2, 42) = 1.28, *p* = 0.29) and local density metrics (F(2, 42) = 0.80, *p* = 0.46).

**Table 1.** Sleep architecture in all experimental groups. Sleep architecture (mean time and standard errors of the means; in minutes) during the 90-min daytime nap for each of the physical practice (n = 15), motor imagery (n = 15) and action observation (n = 15) groups. PP: Physical practice, MI: Motor imagery, AO: Action observation, NREM: Non-Rapid Eye Movement sleep, REM: Rapid Eye Movement sleep.

|  | PP |  | MI |  | AO |  | ANOVAs |  |
| --- | --- | --- | --- | --- | --- | --- | --- | --- |
|  | mean | SEM | mean | SEM | mean | SEM | F | p |
| WAKE (min) | 26.0 | 3.9 | 28.8 | 4.3 | 41.6 | 5.3 | 2.82 | 0.07 |
| NREM1 (min) | 17.5 | 1.8 | 14.0 | 2.0 | 12.2 | 2.0 | 1.98 | 0.15 |
| NREM2 (min) | 24.7 | 2.1 | 29.8 | 3.3 | 25.3 | 3.4 | 1.03 | 0.37 |
| NREM3 (min) | 17.1 | 2.5 | 11.5 | 3.4 | 9.3 | 2.2 | 2.19 | 0.13 |
| REM (min) | 2.6 | 1.6 | 2.4 | 1.2 | 1.0 | 0.6 | 0.54 | 0.59 |

**Table 2.**
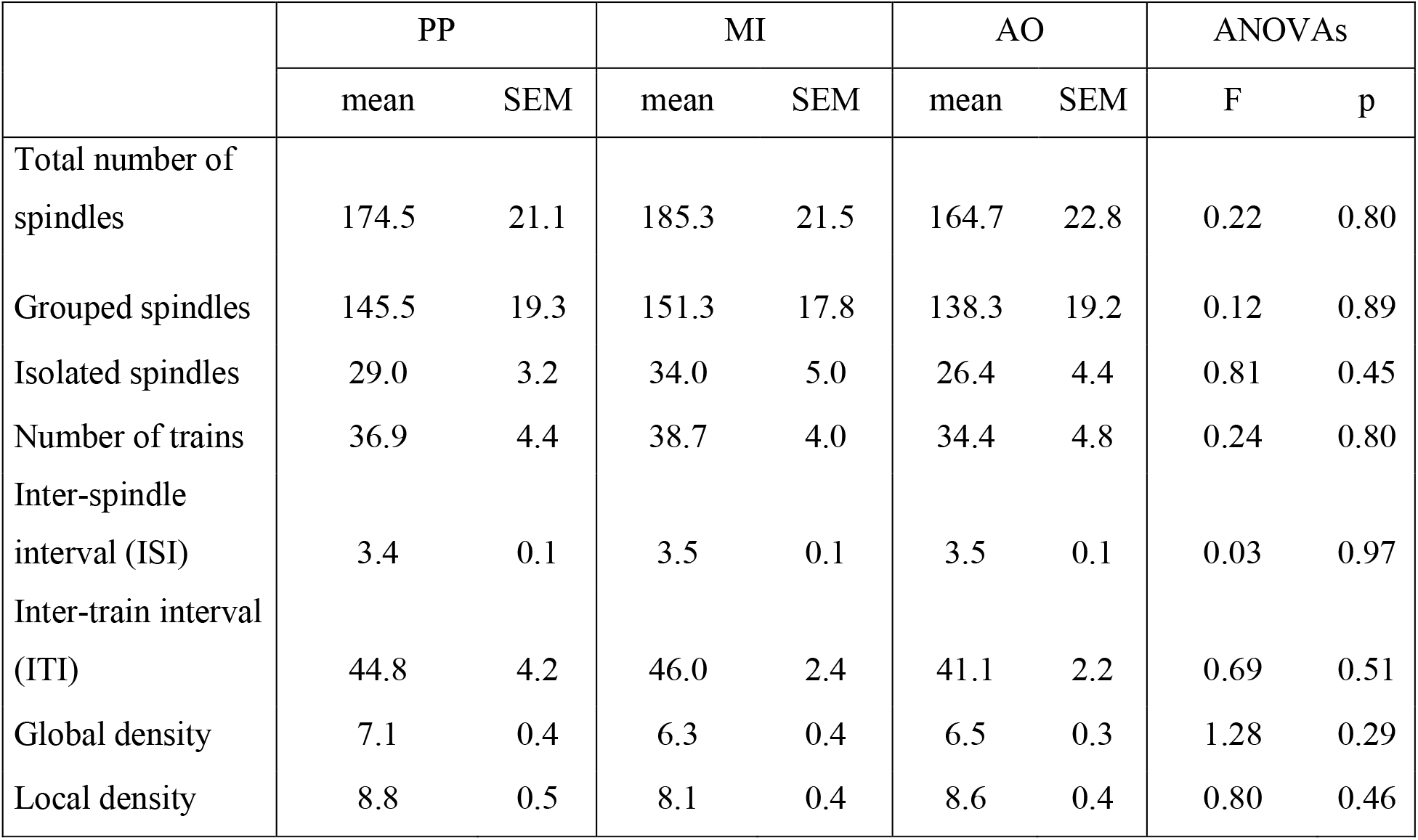
Sleep spindle characteristics in all experimental groups. Means and standard errors of the means (SEM) are reported for the total number of spindles, grouped spindles, isolated spindles, the number of trains, inter-spindle interval (ISI), inter-train interval (ITI), global density (number of spindles per minute), and local density (number of spindles within a spindle-centered 60 s sliding window) of NREM2 sleep spindles at the Pz electrode (midline parietal) during the 90-min daytime nap for each of the physical practice (n = 15), motor imagery (n = 15) and action observation (n = 15) groups. PP: Physical practice, MI: Motor imagery, AO: Action observation.

|  | PP |  | MI |  | AO |  | ANOVAs |  |
| --- | --- | --- | --- | --- | --- | --- | --- | --- |
|  | mean | SEM | mean | SEM | mean | SEM | F | p |
| Total number of spindles | 174.5 | 21.1 | 185.3 | 21.5 | 164.7 | 22.8 | 0.22 | 0.80 |
| Grouped spindles | 145.5 | 19.3 | 151.3 | 17.8 | 138.3 | 19.2 | 0.12 | 0.89 |
| Isolated spindles | 29.0 | 3.2 | 34.0 | 5.0 | 26.4 | 4.4 | 0.81 | 0.45 |
| Number of trains | 36.9 | 4.4 | 38.7 | 4.0 | 34.4 | 4.8 | 0.24 | 0.80 |
| Inter-spindle interval (ISI) | 3.4 | 0.1 | 3.5 | 0.1 | 3.5 | 0.1 | 0.03 | 0.97 |
| Inter-train interval (ITI) | 44.8 | 4.2 | 46.0 | 2.4 | 41.1 | 2.2 | 0.69 | 0.51 |
| Global density | 7.1 | 0.4 | 6.3 | 0.4 | 6.5 | 0.3 | 1.28 | 0.29 |
| Local density | 8.8 | 0.5 | 8.1 | 0.4 | 8.6 | 0.4 | 0.80 | 0.46 |

#### Modality-specific role of NREM2 sleep upon skill consolidation and transfer

To assess the role of NREM2 sleep in motor skill consolidation and generalizability, the duration of NREM2 sleep periods was respectively correlated with the magnitude of skill consolidation and transfer for each group separately. Pearson correlation analyses revealed a significant positive relationship between the duration of NREM2 sleep and the magnitude of post-nap skill consolidation for both the PP group (*r* = 0.53, *p* = 0.041) and the MI group (*r* = 0.62, *p* = 0.015), but not for the AO group (*r* = 0.01, *p* = 0.98). Interestingly, however, a positive relationship between the duration of NREM2 sleep and the magnitude of skill transfer was found in the AO group (*r* = 0.54, *p* = 0.038), but not in the PP (*r* = -0.20, *p* = 0.47) and MI groups (*r* = -0.20, *p* = 0.48), pointing towards modality-specific effects of sleep upon skill consolidation and inter-manual transfer.

#### Relation between sleep spindles and skill consolidation

To investigate the modality-specific contribution of sleep spindle activity in motor skill consolidation, we performed correlation analyses between NREM2 sleep spindle characteristics and the magnitude of skill consolidation for each group separately. For both the PP and MI groups, correlation analyses revealed that the magnitude of skill consolidation correlated positively with the total number of sleep spindles (*r*_PP_ = 0.60, *p* = 0.019; *r*_MI_ = 0.62, *p* = 0.014) (Fig. 3A), with the number of grouped spindles (*r*_PP_ = 0.59, *p* = 0.022; *r*_MI_ = 0.64, *p* = 0.010) (Fig. 4A and 4B), as well as with the number of spindle trains (*r*_PP_ = 0.62, *p* = 0.013; *r*_MI_ = 0.55, *p* = 0.034). In contrast, no significant correlations were found in the AO group between the magnitude of skill consolidation and the total number of sleep spindles (*r* = 0.06, *p* = 0.83) (Fig. 4A), the number of grouped spindles (*r* = 0.09, *p* = 0.76) (Fig. 4C), and the number of spindle trains (*r* = 0.02, *p* = 0.95). It is noteworthy, however, that no significant correlations were found between skill consolidation and the number of isolated spindles for all groups (*r*_PP_ = 0.40, *p* = 0.14; *r*_MI_ = 0.39, *p* = 0.15; *r*_AO_ = -0.06, *p* = 0.83). We performed the Benjamini-Hochberg procedure to control the false discovery rate (FDR)^39^ for the PP and MI groups regarding multiple correlation analyses between skill consolidation and the four spindle metrics (total number of spindles, grouped spindles, spindle trains and isolated spindles). The p-value threshold to reject the null hypothesis after correction is 0.0375 (Ntest = 4) for both groups, confirming the significance of the relationship between the total number of spindles, grouped spindles, and spindle trains with the magnitude of skill consolidation for the PP and MI groups. Considering *r* values of .10, .30, and .50 as the thresholds for small, medium, and large effect sizes, respectively^40^, our correlation analyses are in favor of greater involvement of grouped over isolated spindles in the memory consolidation process following PP and MI. Additional multiple regression analyses can be found in the supplementary material (see Note S4.). Altogether, these findings suggest that sleep spindle activity underlies the consolidation of the practiced motor sequence following PP and MI.

**Fig. 3.**
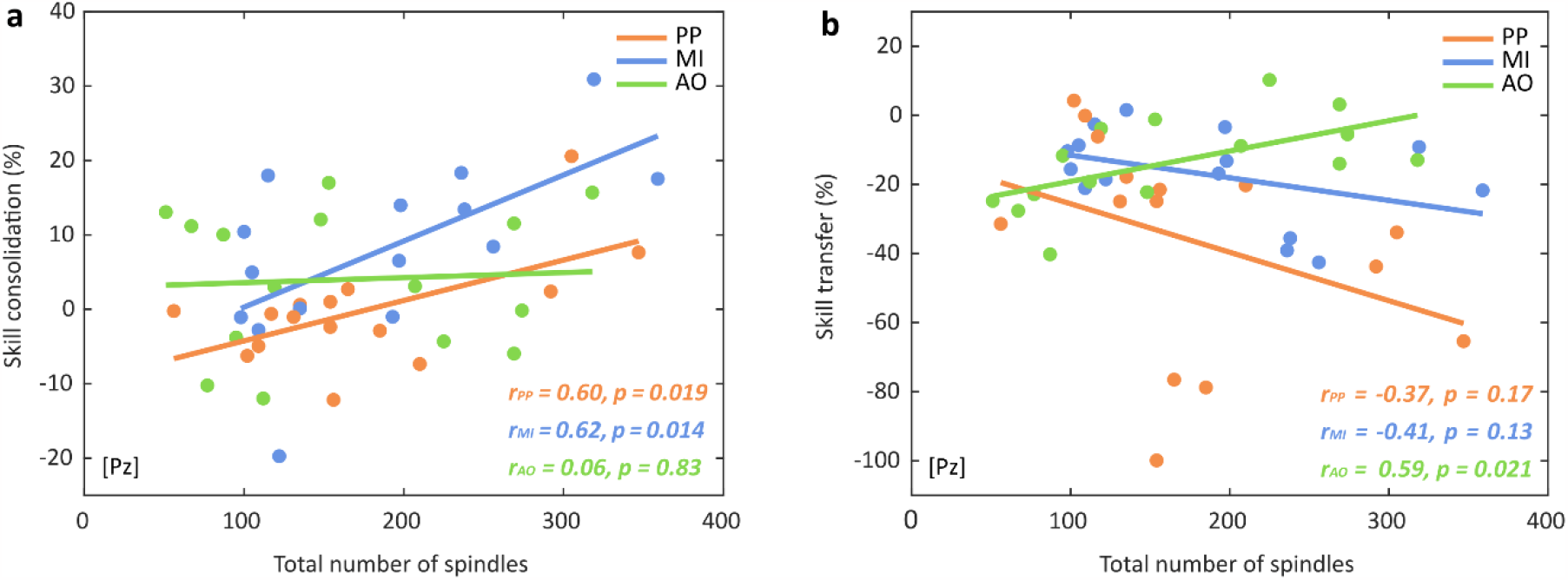
Sleep spindles in relation with motor skill consolidation and transfer. **(a)** Relationship between skill consolidation (percentage of performance changes across the sleep retention interval, from the post-test to the retention test) and the total number of NREM2 spindles for each of the physical practice (PP; n = 15), motor imagery (MI; n = 15) and action observation (AO; n = 15) groups. **(b)** Relationship between skill transfer (percentage of performance changes from the retention test to the inter-manual transfer test) and the total number of NREM2 spindles for each practice group. Scatter plots and linear trend-lines are provided. Pearson correlation coefficients (*r*) and associated p-values are reported for each correlation.

**Fig. 4.**
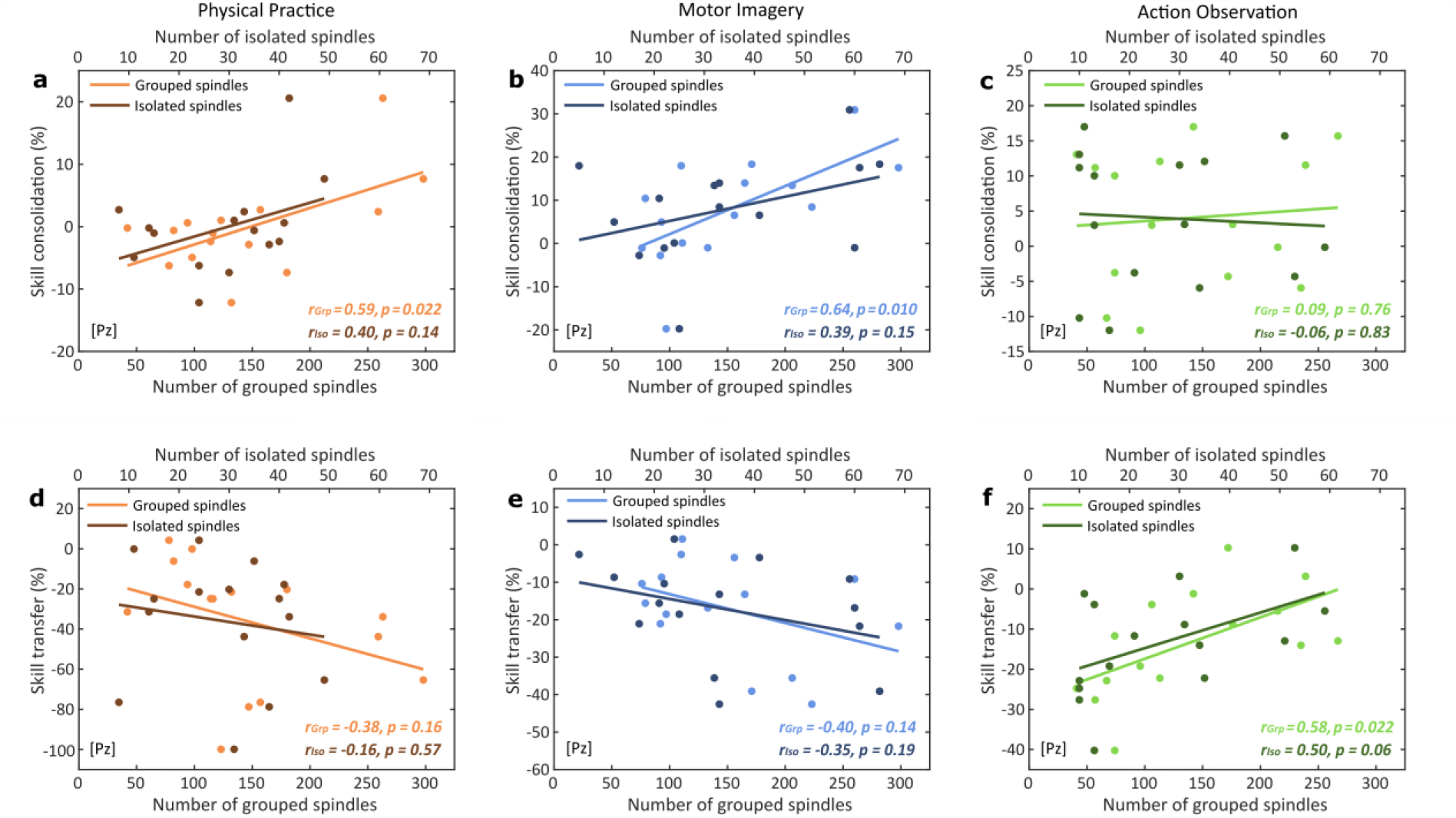
Sleep-related motor skill consolidation and transfer following physical practice, motor imagery and action observation. *Upper row*. Graphical illustration of the relationship between the magnitude of skill consolidation (percentage of performance changes across the sleep retention interval, from the post-test to the retention test) and the total number of grouped (lighter points) or isolated spindles (darker points) in the **(a)** physical practice (red; n = 15), **(b)** motor imagery (blue; n = 15), and **(c)** action observation (green; n = 15) groups. *Lower row*. Graphical illustration of the relationship between the magnitude of skill transfer (percentage of performance changes from the retention to the inter-manual transfer test) and the total number of grouped (lighter points) or isolated spindles (darker points) in the **(d)** physical practice (red; n = 15), **(e)** motor imagery (blue; n = 15), and **(f)** action observation (green; n = 15) groups. Scatter plots and linear trend-lines are provided. Pearson correlation coefficients (*r*) and associated p-values are reported for each correlation.

#### Relation between sleep spindles and skill transfer capacity

To investigate the modality-specific role of sleep spindles in motor skill generalizability, we performed correlation analyses between NREM2 sleep spindle characteristics and the magnitude of skill transfer for each group separately. For the AO group only, significant positive correlations were found between the magnitude of skill transfer and the total number of sleep spindles (*r* = 0.59, *p* = 0.021) (Fig. 3B), the number of grouped spindles (*r* = 0.58, *p* = 0.022) (Fig. 4F), as well as the number of spindle trains (*r* = 0.56, *p* = 0.029). In contrast, no significant correlations were found in the PP and MI groups between the magnitude of skill transfer and the total number of sleep spindles (*r*_PP_ = -0.37, *p* = 0.17; *r*_MI_ = -0.41, *p* = 0.13) (Fig. 3B), the number of grouped spindles (*r*_PP_ = - 0.38, *p* = 0.16; *r*_MI_ = -0.40, *p* = 0.14) (Fig. 4D and 4E), and the number of spindle trains (*r*_PP_ = - 0.29, *p* = 0.30; *r*_MI_ = -0. 31, *p* = 0.26). For all groups, no significant correlations were found between the magnitude of skill transfer and the number of isolated spindles (*r*_PP_ = -0.16, *p* = 0.57; *r*_MI_ = -0.35, *p* = 0.19; *r*_AO_ = 0.50, *p* = 0.06). Again, we performed the Benjamini-Hochberg procedure to control the false discovery rate^39^ for the AO group regarding multiple correlation analyses between skill transfer and the four spindle metrics (total amount of spindles, grouped spindles, spindle trains and isolated spindles). The p-value threshold to reject the null hypothesis after correction is 0.0375 (Ntest = 4), confirming the significance of the relationship between the total number of spindles, grouped spindles and spindle trains with the magnitude of skill transfer for the AO group. Considering the effect sizes of the correlations^40^, our findings are slightly in favor of greater involvement of grouped over isolated spindles in motor skill transfer capacity following AO. Additional multiple regression analyses can be found in the supplementary material (see Note S4.). Altogether, these findings suggest that sleep spindle activity promotes the consolidation of an effector-unspecific representation (i.e., transfer to the unpracticed hand) of the learned motor sequence following AO.

#### Time-frequency maps

Fig. 5 depicts the grand average time-frequency (TF) maps zoomed in on -6 s and +6 s around spindle onsets during NREM2 sleep for the PP (left panel), MI (middle panel) and AO (right panel) groups. To assess within-group power variations at each frequency bin over time, we used a two-tailed student’s *t*-test against a predefined baseline window (−2 to -0.5 seconds before spindle onsets) for each frequency bin. TF maps were then FDR-corrected using the Benjamini-Hochberg procedure^39^ (Ntest = 60020). TF analyses confirmed the clustering and rhythmic nature of spindle events in all groups. The group-averaged TF maps illustrate the significant periodic power increases in the spindle frequency band every 3-4 seconds (∼0.2-0.3 Hz), irrespective of the practice mode. We also conducted independent sample student’s t-tests to evaluate between-groups differences of the spectral power variations. Similarly, TF maps were corrected for multiple comparisons using the Benjamini-Hochberg procedure (Ntest = 60020). No significant difference was found between the groups, underlying the modality-unspecific pattern of the spindle-power rhythmic variations over time. Noteworthy, TF maps also reveal a rhythmic pattern of power increases in the theta frequency band (4-8 Hz), which accords with current trends suggesting that cross-frequency interactions between sleep spindles and theta waves may be relevant for sleep-related memory consolidation (see also ^41^).

**Fig. 5.**
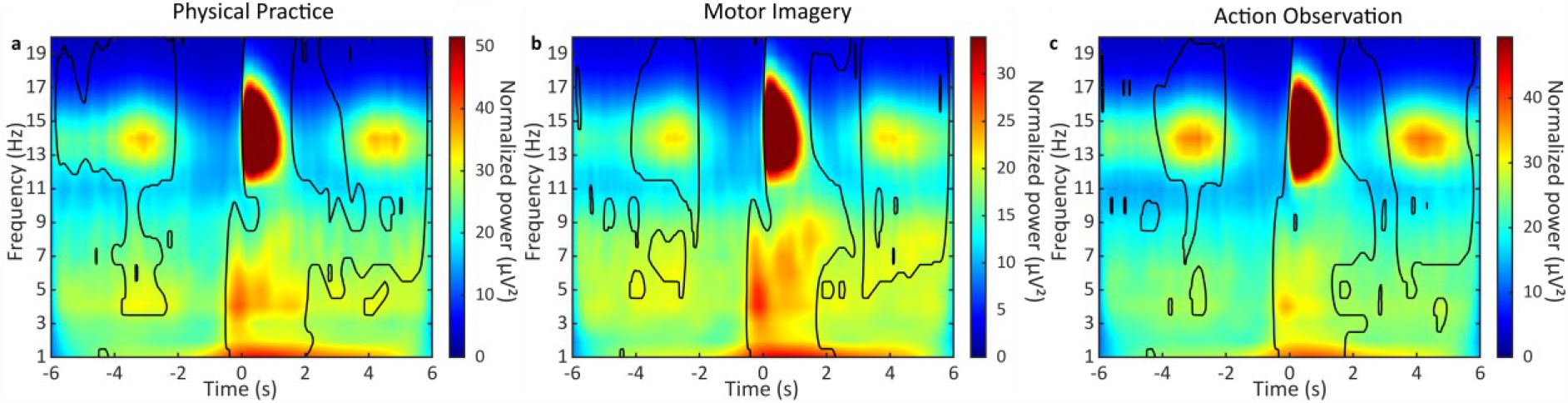
Time-frequency decomposition of NREM2 sleep spindles following physical practice, motor imagery and action observation. Grand average time-frequency maps across participants for spindle events occurring at scalp electrode Pz, using epoch windows ranging from -6 s to +6 s around spindle onsets, and illustrating the 0.2-0.3 Hz spindle rhythmicity within trains during NREM2 sleep periods following physical practice (left panel; n = 15), motor imagery (middle panel; n = 15) and action observation (right panel; n = 15). The color bar reflects normalized (1/f compensation) spectral power values (µV²). Contour lines indicate regions where power is significantly higher than the baseline (−2 s to -0.5 s) for each frequency bin after correction for multiple comparisons using the Benjamini-Hochberg procedure to control the false discovery rate (N_test_ = 60020). NREM: Non-Rapid Eye Movement.

## Discussion

In the current study, we examined (i) whether similar consolidation mechanisms are engaged during sleep following PP, MI and AO practice, and (ii) the contribution of sleep spindles in motor skill consolidation and transfer, depending on the modality of practice. Our findings confirmed that participants acquired the motor sequence through PP, MI and AO practice, with an advantage for PP. Our results further revealed that sleep, and more specifically the time spent in NREM2 sleep, is related to motor skill consolidation following PP and MI practice and motor skill transfer following AO practice, hence pointing towards potential modality-specific effects of sleep upon skill consolidation and its generalizability. In addition, we found that spindles occurring in trains (grouped spindles) during NREM2 sleep are primarily involved in the sleep consolidation process, in comparison to isolated ones, leading to enhanced skill retention following PP and MI practice and improved skill transfer following AO practice. Finally, our results revealed that irrespective of the modality of practice, spindles tend to cluster in trains on a low-frequency time scale of about 50 s (∼0.02 Hz), and during which spindles iterate every 3-4 s (∼0.2-0.3 Hz). Therefore, and for the first time, our results provide evidence of a modality-unspecific organization of sleep spindles during the consolidation of motor skills. During training, and as expected, the rehearsal of the motor sequence increased performance for all practice groups^4–6,13,16,42–44^. As expected, though, participants in the PP group expressed higher performance improvements than their MI group counterparts^13,16,42,45^. Traditionally, it is also assumed that PP leads to greater practice-related gains than AO^5,14,15^. Albeit not significant, a clear tendency emerged in our results in accordance with this latter assumption. Previous studies have shown that AO learners may reach similar performance levels than PP learners when only a few physical practice trials are provided^5,15^. Here, we can thus assume that motor-related information during AO may have been partly obtained through actual execution during the pre-test, leading to the encoding of a sequence representation calibrated with prior physical execution and not only because of additional physical practice during the test blocks since similar results would have emerged in the MI practice condition.

In relation to motor skill consolidation, our results first corroborated previous findings in showing significant over-nap performance gains for all groups (see Fig S1.)^32,34,46–48^. To better examine the modality-specific effects of sleep upon motor skill consolidation, we evaluated the relationship between the magnitude of post-nap performance changes and sleep characteristics in the PP, MI and AO groups. We found a positive relationship between the time spent in NREM2 sleep and the magnitude of motor skill consolidation for participants in the PP and MI groups. This finding accords with previous studies depicting the crucial role of NREM2 sleep in motor memory consolidation following PP^25,32,34,48^. To the best of our knowledge, this is the first demonstration of the role of NREM2 sleep in the consolidation of motor skills following MI practice, albeit theoretically assumed^11,51^. In addition, we also found a positive relationship between the number of NREM2 sleep spindles and the magnitude of motor skill consolidation following PP and MI practice, emphasizing the role of sleep spindles in motor memory consolidation, even in the absence of overt movement. In contrast, however, our findings did not reveal any relationship between NREM2 sleep or spindle activity with the magnitude of skill consolidation following AO practice. These results are, at first sight, difficult to reconcile with the seminal study of Van Der Werf et al. (2009)^11^, who demonstrated that an early sleep window following AO is crucial for the emergence of offline performance gains. However, in their study, the benefits of sleep were only observed in behavioral performance at the group level, and the absence of polysomnographic monitoring prevented any analysis of sleep architecture or spindle activity in relation to motor skill consolidation. Thus, our results rather accord with studies suggesting that AO practice may trigger different consolidation processes than those triggered by PP, leading to different behavioral outcomes^14,52^.

In this way, and very interestingly, our results confirmed the potential engagement of distinct consolidation mechanisms during sleep following PP, MI and AO. Indeed, our findings revealed a positive relationship between the time spent in NREM2 sleep and inter-manual skill transfer for participants in the AO group only. We further found a positive correlation between the number of NREM2 sleep spindles and the magnitude of skill transfer following AO practice, revealing the role of sleep spindle activity in the memory consolidation process following AO. Hence, we conjecture that sleep differently affects the representation of a motor skill depending on prior training experience. This assumption is also supported by transfer performance at the group level since we showed greater post-nap skill transfer following both MI and AO compared to PP, which accords with previous studies^5,17,18,20,45^. It is now well accepted in the literature that PP of long complex sequences would mainly engage an encoding of the motor task in visuo-spatial coordinates (i.e., effector-unspecific learning)^2,5,53^. In contrast, shorter sequences would mainly be encoded in motor coordinates (i.e., effector-specific learning)^2,54,55^. Therefore, physical practice of a short 5-element motor sequence in our study may have primarily led to the development of an effector-specific learning of the motor skill, as reflected by impaired transfer performance compared to other practice modalities. In contrast, the higher inter-manual skill transfer capacity found for the MI and AO groups indicates that both non-physical practice conditions may have led to the development of a motor skill representation mostly at an effector-unspecific level, in comparison to the PP group (see also ^5,45^).

Our behavioral results also accord with previous neuroimaging findings showing more consistent brain activations during MI and AO compared to PP, in an effector-unspecific manner within a predominant premotor-parietal network^8,9,56,57^. In a recent neuroimaging meta-analysis, Hardwick and colleagues^8^ compared the pattern of brain activations following PP, MI and AO, and identified brain areas involved in both the simulation of actions (MI and AO) and actual motor execution (PP) through a cortical-dominant network. This network comprises essentially premotor, parietal, and sensorimotor regions, albeit to a lesser extent for MI and AO. Hence, while a set of common neural structures is activated during PP, MI and AO, leading to enhanced motor performance and learning, the additional and specific recruitment of brain regions with respect to the modality of practice may be responsible for the distinct encoding strategies and behavioral outcomes observed during the post-training, retention and transfer tests.

Finally, we investigated the organization of sleep spindle activity in all practice groups, and its potential contribution to motor skill consolidation and transfer. Interestingly, time-frequency analyses revealed an apparent clustering and temporal occurrence of sleep spindles every 3 to 4 seconds in all practice groups. Hence, such a cluster-based organization of sleep spindles in trains may be an endogenous, modality-unspecific mechanism critical for consolidating newly-formed motor memories during sleep. In addition, and as previously shown for PP-based motor learning^36,37,58,59^, our analyses confirm that spindles tend to cluster on an infraslow time scale of about 50 s (∼0.02 Hz) during NREM2 sleep, with a rhythmic occurrence of spindles during trains that follows a time scale of about 3-4 s (∼0.2-0.3 Hz), irrespective of the practice mode. In order to assess the relevance of this temporal organization, sleep spindles were categorized as grouped or isolated according to whether or not they belonged to spindle trains, respectively (see STAR Methods). Correlation analyses revealed significant positive relationships between grouped spindles and the magnitude of skill consolidation following PP and MI, but not AO. In contrast, a significant positive relationship was found between grouped spindles and the magnitude of skill transfer in the AO practice group only. No significant correlations were found between isolated spindles and the magnitude of skill consolidation and transfer for all groups. These results extend recent findings regarding overt and covert practice by showing that grouped spindles seem more involved than isolated ones in the consolidation and effector transfer capacity of motor skills acquired through physical and non-physical practice^11,25,27,31,37,52^. Therefore, despite an apparent modality-unspecific and cluster-based organization of NREM2 sleep spindles during the post-learning nap, we conjecture that spindles may be involved in the sleep consolidation of motor skills through the reactivation of modality-specific brain regions, leading to the distinct behavioral outcomes observed during retention and inter-manual transfer performance.

It should be noted, however, that we failed to find significant relationships between the magnitude of skill consolidation and transfer with both global and local spindle density metrics, which are commonly used in the memory consolidation and sleep research domains^31^. This is likely due to a difference in spindle activity between a nap and a night of sleep. Indeed, substantial variations in the sleep EEG spectra (in the spindle frequency band in particular) that depend on melatonin rhythms and endogenous circadian phases of sleep consolidation^60,61^ could explain the absence of significant correlations between spindle density metrics and post-nap performance in our study. A density calculation over such a short sleep time may therefore not be adequate to quantify the direct relationship between sleep spindles and motor memory consolidation (see ^34^).

To conclude, physical practice, motor imagery and action observation are effective practice modalities to learn a motor skill, with the development of an effective memory representation during practice that undergoes further modification during subsequent sleep. Interestingly, our findings underline the modality-specific role of NREM2 sleep spindle activity in motor skill consolidation and transfer. Despite an apparent similar cluster-based organization of NREM2 spindles during the post-learning nap in all groups, different behavioral outcomes are elicited during retention and transfer performance. We show that a daytime nap offers an early sleep window that promotes the consolidation and retention of the learned motor skill following PP and MI practice, while an effector-unspecific sequence representation favoring skill transfer is consolidated after AO practice. Altogether, we demonstrate that PP per se is not a pre-requisite for sleep-related consolidation of motor skills and that the clustering of sleep spindles in trains may be an inherent memory-reprocessing mechanism for the effective consolidation of modality-specific skill representations. Finally, given the non-physical nature of MI and AO practice, and their sleep-related skill consolidation and transfer opportunities, our findings may encourage the development of non-physical training protocols for the learning of new motor skills, as well as for the designing of innovative rehabilitation interventions such as for patients with motor deficits having to remaster skills following physical or brain injury.

## Limitations of the study

In this study, we used a nap paradigm and recruited exclusively young healthy participants. It would be interesting then to study the potential involvement and temporal organization of sleep spindles in the consolidation of motor skills learned by PP, MI and AO practice during a whole night of sleep or with clinical populations facing spindle abnormalities or altered rhythmicity of spindles, for instance.

We conducted our main analyses on the parietal Pz derivation of the EEG cap, described as sensitive to sleep spindle activity in relation to motor memory consolidation^31,32,62^. Relationships with other derivations are described in the supplemental material (see Fig. S2 and S3). However, further analyses may be relevant to assess and precise the spatial dynamics of the cluster-based organization of sleep spindles over different brain regions.

Regarding the evaluation of motor performance, we decided to administer only a single block of practice during the retention and transfer tests, leaving little room for additional physical practice. It was necessary in our study to limit the physical execution of the sequence to prevent additional training during the test phases, and especially for the non-physical MI and AO practice groups. However, such a design implies reduced measures of RT performance and therefore greater performance variability compared to more common designs using two or more blocks during testing phases^31,49^.

Finally, while our correlation analyses are in favor of greater involvement of grouped over isolated spindles in the sleep-related memory consolidation process, additional statistical comparisons of the coefficient correlations between groups (by conducting Fisher’s r to z transform) failed to reveal any significant group difference. However, it must be noted that a large sample size (N = 177) is necessary to detect a medium-sized difference of two Pearson r coefficients (Δr ∼ 0.3)^40^.

## Author Contributions

Author contributions: A.C., A.B. and U.D. conceived the experiment; A.C. collected the data; A.C. and A.B. analyzed the data and discussed the results; A.C., A.B. and U.D. wrote the manuscript; All authors revised the manuscript.

## Material and methods

### RESSOURCES AVAILABILITY

#### Lead contact

Further information should be directed to and will be fulfilled by the lead contact, Arnaud Boutin

#### Data and code availability

The data that support the results of this study are available from the corresponding author upon reasonable request and under a formal data-sharing agreement.

Sleep EEG data were processed using the MATLAB R2019b software from The MathWorks (Natick, MA) and the open-source Brainstorm and EEGLAB software. The codes for the detection and clustering of sleep spindles are available at the following GitHub repositories: https://github.com/arnaudboutin/Spindle-detection and https://github.com/arnaudboutin/Spindle-clustering. The codes used to perform other analyses are available from the corresponding author upon reasonable request.

Any additional information required to reanalyze the data reported in this paper is available from the lead contact upon request.

### EXPERIMENTAL MODEL AND SUBJECT DETAILS

Forty-five healthy volunteers (18 females, mean age: 23.7 ± 4 years) were recruited by local advertisements and were randomly and equally assigned to either a PP group (8 females, mean age: 22.8 ± 4 years), MI group (6 females, mean age: 24.5 ± 4 years) or AO group (4 females, mean age: 23.9 ± 5 years). All participants met the following inclusion criteria: aged between 18 and 35 years, right-handed (Edinburgh Handedness Inventory^63^), medication-free, without history of mental illness, epilepsy or head injury with loss of consciousness, sleep or neurologic disorders and no recent upper extremity injuries. The Local Ethics Committee from the Université Paris-Saclay (CER-Paris-Saclay-2019-057-A1) approved the experimental protocol, which conformed to relevant guidelines and regulations. All participants gave written informed consent before inclusion. Participants were asked to maintain a regular sleep-wake cycle and refrain from all caffeine- and alcohol-containing beverages 24h before the experimentation. Participants in the MI group were also required to complete the French version of the Movement Imagery Questionnaire-Third version (MIQ-3f)^64^ before starting the experimental protocol. The MIQ-3f is a twelve-item self-report questionnaire, in which participants are asked to perform a given movement followed by its mental execution either by external visual imagery, internal visual imagery or kinesthetic imagery. Participants rate the difficulty with two 7-point scales respectively for visual or kinesthetic imagery, ranging from 1 (very hard) to 7 (very easy). A higher average score represents a greater imagery capacity.

## METHOD DETAILS

### Experimental design and motor sequence task

Participants sat on a chair at a distance of 50 cm in front of a computer screen, equipped with a 64-channel EEG cap. The motor task consisted of performing as quickly and accurately as possible a 5-element finger movement sequence by pressing the appropriate response keys on a standard French AZERTY keyboard using their left, non-dominant hand. The sequence to be performed (C-N-V-B-C, where C corresponds to the little finger and N to the index finger) was explicitly taught to the participant prior to training.

Physical practice blocks consisted of 16 repetitions of the 5-element sequence (i.e., a total of 80 keypresses). Each block began with the presentation of a green cross in the center of the screen accompanied by a brief 50-ms tone. In case of occasional errors, participants were asked “not to correct errors and to continue the task from the beginning of the sequence” (see ^65^ for a similar procedure). At the end of each block, upon completion of the 80 keypresses, the color of the green-colored imperative stimulus turned red, and participants were then required to look at the fixation cross during the 30-s rest period. This protocol controlled the number of movements executed per block to ensure that the same amount of practice with the task was afforded to participants during a particular session. Stimuli presentation and response registration were controlled using the MATLAB R2016b software from The MathWorks (Natick, MA) and the Psychophysics Toolbox extensions^66^.

The study started at 1.00 pm to minimize the putative impact of both circadian and homeostatic factors on individual performance levels and sleep characteristics^60,61,67^. The experimental procedure was composed of seven main phases: familiarization, pre-test, training, post-test, 90-minute nap, retention and transfer tests (Fig. 1). Before training, participants underwent a brief familiarization phase during which they were instructed to repeatedly and slowly perform the 5-element sequence until they accurately reproduced the sequence three consecutive times. This familiarization was intended to ensure that participants understood the instructions and explicitly memorized the sequence of movements.

During the pre-test, all participants physically performed one block of the 5-element motor sequence. The ensuing training phase consisted of 14 blocks performed with physical, observational or mental practice. Participants in the PP group were asked to physically execute the sequence task with their left-hand fingers, as previously described. Participants in the AO group were instructed to keep their fingers on the corresponding response keys. Following the imperative green-cross stimulus and audio cue, a video of a model performing the motor task was displayed on the screen. The model was depicted so that the observers could see both the finger movements of the model and the green cross appearance on the screen. This viewing angle was adopted in order to closely match the perspective view of the AO participants. An additional window inset zooming on the left-hand fingers of the model was implemented so that participants could precisely watch fine finger movements. Participants in the AO group were free to observe both perspectives in an active and conscious manner while avoiding any concurrent muscular execution of the movement (controlled online by electromyography (EMG) recording electrodes placed on the left flexor digitorum superficialis; see section EEG-EMG data acquisition and pre-processing for details). Based on previous motor sequence learning findings^31^, performance improvements of the model across training blocks followed the power-law of practice (mean response time between consecutive keypresses ranging from RT_Block1_ = 616 ms to RT_Block14_ = 223 ms). Participants in the MI group were instructed to keep their left-hand fingers on the corresponding response keys and their thumb on the keyboard’s space bar. When they heard the imperative audio cue, they had to imagine themselves performing the sequence using a combination of internal visual and kinesthetic imagery, while avoiding any associated overt movements (controlled online by similar EMG procedure as for the AO group). After completing each mentally rehearsed sequence, they were asked to press the space bar with their thumb to objectively control for the amount of MI practice. Each block was composed of 16 mental repetitions of the motor sequence.

Approximately five minutes after completion of the training phase, all participants underwent a post-test phase identical to the pre-test. This session was briefly preceded by a physical warm-up phase (i.e., slow-paced production of the sequence three consecutive times) for all groups. Following the post-test, all participants were administered a 90-minute nap. Ten minutes after awakening, all participants were asked to perform again a physical warm-up phase (i.e., three slow-paced repetitions of the trained sequence) before completing the retention and transfer test blocks. The retention test was done first and consisted of one practice block performed with the non-dominant left hand. An inter-manual transfer test was then carried out. Participants were instructed to perform one block on the original motor sequence but with the unpracticed, dominant right hand.

### EEG-EMG data acquisition and pre-processing

#### EMG recordings

Two bipolar Ag-AgCl electrodes with 10 kΩ safety resistor were placed at a distance of 3 cm from each other along the belly of the left flexor digitorum superficialis. During training, the EMG signal was monitored and controlled online to ensure the absence of micro finger movements for the MI and AO groups. If the experimenter detected any muscle activity, participants were instructed to immediately relax their hand.

#### EEG recordings

EEG was acquired using a 64-channel EEG cap (actiCAP snap BrainProducts Inc.). The EEG cap included slim-type electrodes (5kΩ safety resistor) suitable for sleep recordings, with FCz and AFz being, respectively, the reference and ground electrodes during the recordings. For reliable sleep stage scoring, we also added electrooculography (EOG) and EMG recordings using bipolar Ag-AgCl electrodes. We recorded the vertical EOG component by placing pairs of electrodes above and below the left eye. EMG bipolar electrodes were placed over the chin. All EEG, EMG and EOG data were recorded using two battery-powered 32-channel amplifiers and a 16-channel bipolar amplifier (respectively, BrainAmp and BrainAmp ExG, Brain Products Inc.). All signals were recorded at a 5-kHz sampling rate with a 100-nV resolution. Electrode-skin impedance was kept below 5 kΩ using Abralyt HiCl electrode paste to ensure stable recordings throughout all experimental phases.

EEG data were bandpass filtered between 0.5 and 50 Hz to remove low-frequency drift and high-frequency noise, down-sampled to 250 Hz, and re-referenced to the linked mastoids (i.e., TP9 and TP10). EOG and EMG data were bandpass filtered between 0.3-35 Hz and 10-100 Hz, respectively.

## QUANTIFICATION AND STATISTICAL ANALYSIS

All statistical analyses used every participant in each experimental group (PP: n = 15; MI: n = 15; AO: n = 15). These sample sizes were determined based on previous studies^32,36,49^. All error measurements indicate the standard error of the means (SEM).

### Behavioral analysis

RT performance was measured as the interval between two consecutive keypresses during each test block. Also, since participants were asked to start over from the beginning of the sequence if they made any error during task production, RTs from error trials (i.e., erroneous key presses) were excluded from the analyses. Across all test blocks, the PP group performed 14.6 (± 1.6) accurate sequences, 14.1 (± 1.7) for the MI group, and 14.6 (± 1.7) for the AO group (out of a maximum of 16 sequences). To better reflect individual performance on the motor sequence task, we computed mean RT performance on accurately typed sequences^68,69^.

Individual RTs were then averaged to obtain an overall estimation of the performance for each practice block. Motor skill acquisition was assessed by analyzing the RT performance changes (in percentages) from the pre-test to the post-test block. Motor skill consolidation was assessed by analyzing the post-nap retention performance changes, as indexed by the percentage of RT changes from the post-test to the retention test. Finally, inter-manual skill transfer capacity was assessed by computing the difference (in percentages) between RT performance on the retention and transfer test blocks. Note that negative values reflect impairments in skill transfer.

A one-way ANOVA with the between-subject factor MODALITY (PP, MI, AO) was first performed on the RT data during the pre-test to ensure that the three groups did not differ at baseline performance. Then, one-way ANOVAs were applied separately for motor skill acquisition, consolidation and transfer. The significance threshold was set at 0.05 for all analyses. Holm post-hoc comparisons were performed in case of significant effects or interactions.

### EEG analysis

#### Spindle detection

The artifact-free EEG signal was sleep-stage scored according to AASM guidelines^70^. Each 30-second epoch was visually scored as either NREM stages 1-3, REM, or wake (Table 1). The detection of spindle events was conducted using all artifact-free NREM2 sleep epochs over the parietal site (electrode Pz), as expression of spindles has been shown to predominate over this region following motor sequence learning^31,32,37,62^. Discrete sleep spindle events (i.e., onset and offset) were automatically detected using a wavelet-based algorithm (see ^31^ for further details). Spindles were detected at the Pz derivation by applying a dynamic thresholding algorithm (based on the median absolute deviation of the power spectrum) to the extracted wavelet scale corresponding to the 11-16 Hz frequency range and a minimum window duration set at 300 ms^31,37,71^ (see ^25^ for a review). Events were considered sleep spindles only if they lasted 0.3-2 seconds, occurred within the 11-16 Hz frequency range and with onsets during NREM2 sleep periods.

#### Spindle clustering

As recently proposed by Boutin and Doyon (2020)^25^, sleep spindles may be split into two categories: clusters of two or more consecutive and electrode-specific spindle events interspaced by less than or equal to 6 seconds were operationalized as trains, in comparison to those occurring in isolation (i.e., more than 6 seconds between two consecutive spindles detected on the same electrode). Hence, for convenience, spindles belonging to trains were categorized as *grouped spindles*, and those occurring in isolation were categorized as *isolated spindles*. Several variables of interest were considered: the total number of spindles, three metrics related to spindle clustering (number of grouped and isolated spindles, number of trains), two measures of spindle rhythmicity (inter-spindle interval [ISI] within trains and inter-train interval [ITI]; in seconds), and two measures of spindle density (global density [number of spindles per minute] and local density [number of spindles within a spindle-centered sliding window of 60 seconds]). One-way ANOVAs with the between-subject factor MODALITY (PP, MI, AO) were applied separately for each spindle metric. In addition, Pearson correlation analyses were performed to evaluate the relationship between sleep spindles and the magnitude of skill consolidation and transfer. The significance threshold was set at 0.05 for all analyses. If significant correlations were found, we used the Benjamini-Hochberg procedure to adjust the p-value and control the false discovery rate^39^.

#### Time-frequency maps

To evaluate the temporal organization of NREM2 sleep spindles, we conducted time-frequency analyses. For each NREM2 sleep spindle detected, we extracted a 12-second time window centered on the spindle onset. We applied for each epoch a TF decomposition across the 1-20 Hz frequency range using complex Morlet Wavelet. Spectral resolution of the wavelet was defined with a central frequency set at 1 Hz, and temporally with the full-width at half-maximum (FWHM) set at 3 seconds. For illustration purposes, TF maps were normalized by multiplying the power at each frequency bin with the frequency value (1/f compensation). To assess the statistical significance of the power variation at each frequency bin over time, we used a two-tailed student’s *t*-test against a predefined baseline window for each practice group with a significance threshold set at 0.05. The baseline was set from -2 s to -0.5 s before spindle onset. We also conducted two-tailed independent sample student’s t-tests to evaluate between-group differences of the power variation over time. All statistical maps were then corrected for multiple comparisons using the Benjamini-Hochberg procedure to control the false discovery rate (Ntest = 60020 for each analysis)^39^. All analyses were done using the MATLAB R2019b software from The MathWorks (Natick, MA) and the open-source Brainstorm software^72^.

## Supplemental information

**Figure S1. Behavioral results**. Illustration of the RT performance of all groups for each test block. To assess changes in performance at the end of the training phase depending on the initial practice mode, we performed a mixed ANOVA using a MODALITY (PP, MI, AO) x BLOCK (pre-test, post-test) factorial design, with repeated measures on the factor BLOCK. The analysis revealed significant main effects of MODALITY (F(2,42) = 5.25, *p* = 0.009, *ɳ* ^2^_p_ *=* .20) and BLOCK (F(1,42) = 128, *p* < 0.001, *ɳ* ^2^_p_ *=* .75), but failed to detect a significant MODALITY X BLOCK INTERACTION (F(2,42) = 2.02, *p* = 0.15). For the main effect of MODALITY, Holm post-hoc comparisons revealed that participants in the PP group (RT = 341 ms) outperformed participants in both the MI (RT = 425 ms, *p* = 0.017) and AO groups (RT = 419 ms, *p* = 0.020). For the main effect of BLOCK, post-hoc comparisons revealed lower RTs during the post-test relative to the pre-test (from RT_pretest_ = 450 ms to RT_posttest_ = 339 ms, *p <* 0.001).

In addition, to assess sleep-dependent changes in performance depending on the initial practice mode, we performed a mixed ANOVA using a MODALITY (PP, MI, AO) x BLOCK (post-test, retention) factorial design, with repeated measures on the factor BLOCK. The analysis revealed significant main effects of MODALITY (F(2,42) = 8.85, *p* < 0.001, *ɳ* ^2^_p_ *=* .30) and BLOCK (F(1,42) = 9.18, *p* = 0.004, *ɳ* ^2^_p_ *=* .14), but failed to detect a significant MODALITY X BLOCK INTERACTION (F(2,42) = 2.76, *p* = 0.075). For the main effect of MODALITY, Holm post-hoc comparisons revealed that participants in the PP group (RT = 273 ms) outperformed participants in both the MI (RT = 367 ms, *p* = 0.001) and AO groups (RT = 354 ms, *p* = 0.003). For the main effect of BLOCK, post-hoc comparisons revealed lower RTs during the retention test relative to the post-test (from RT_posttest_ = 339 ms to RT_retention_ = 322 ms, *p =* 0.004).

Finally, to assess the inter-manual transfer capacity depending on the initial practice mode, we performed a mixed ANOVA using a MODALITY (PP, MI, AO) x BLOCK (retention, transfer) factorial design, with repeated measures on the factor BLOCK. The analysis revealed significant main effects of MODALITY (F(2,42) = 3.54, *p* = 0.038, *ɳ* ^2^_p_ *=* .14) and BLOCK (F(1,42) = 61.0, *p <* 0.001, *ɳ* ^2^_p_ *=* .59), but failed to detect a significant MODALITY X BLOCK INTERACTION (F(2,42) = 2.18, *p* = 0.13). For the main effect of MODALITY, Holm post-hoc comparisons revealed that participants in the PP group (RT = 316 ms) outperformed participants in the MI group (RT = 380 ms, *p* = 0.049) but not in the AO group (RT = 368 ms, *p* = 0.097). For the main effect of BLOCK, post-hoc comparisons revealed higher RTs during the transfer test relative to the retention test (from RT_retention_ = 323 ms to RT_transfer_ = 387 ms, *p <* 0.001).

**Fig. S1.**
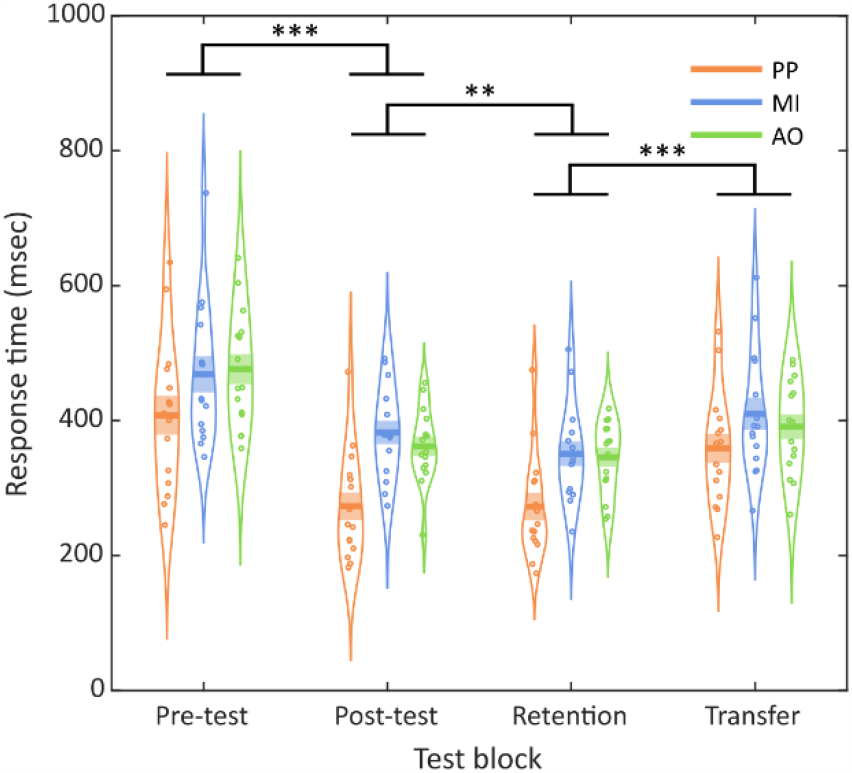
Behavioral results. Mean response time (RT) on each test block according to the modality of practice in the physical practice (PP), motor imagery (MI) and action observation (AO) groups. The curved lines indicate the distribution of data, the dark bars represent the mean of the distribution, and the lighter area surrounding the mean represent the standard error of the means. Individual data points are displayed as colored circles (n = 15 for each group at each test block). ** *p* < 0.01, *** *p* < 0.001.

**Fig. S2.**
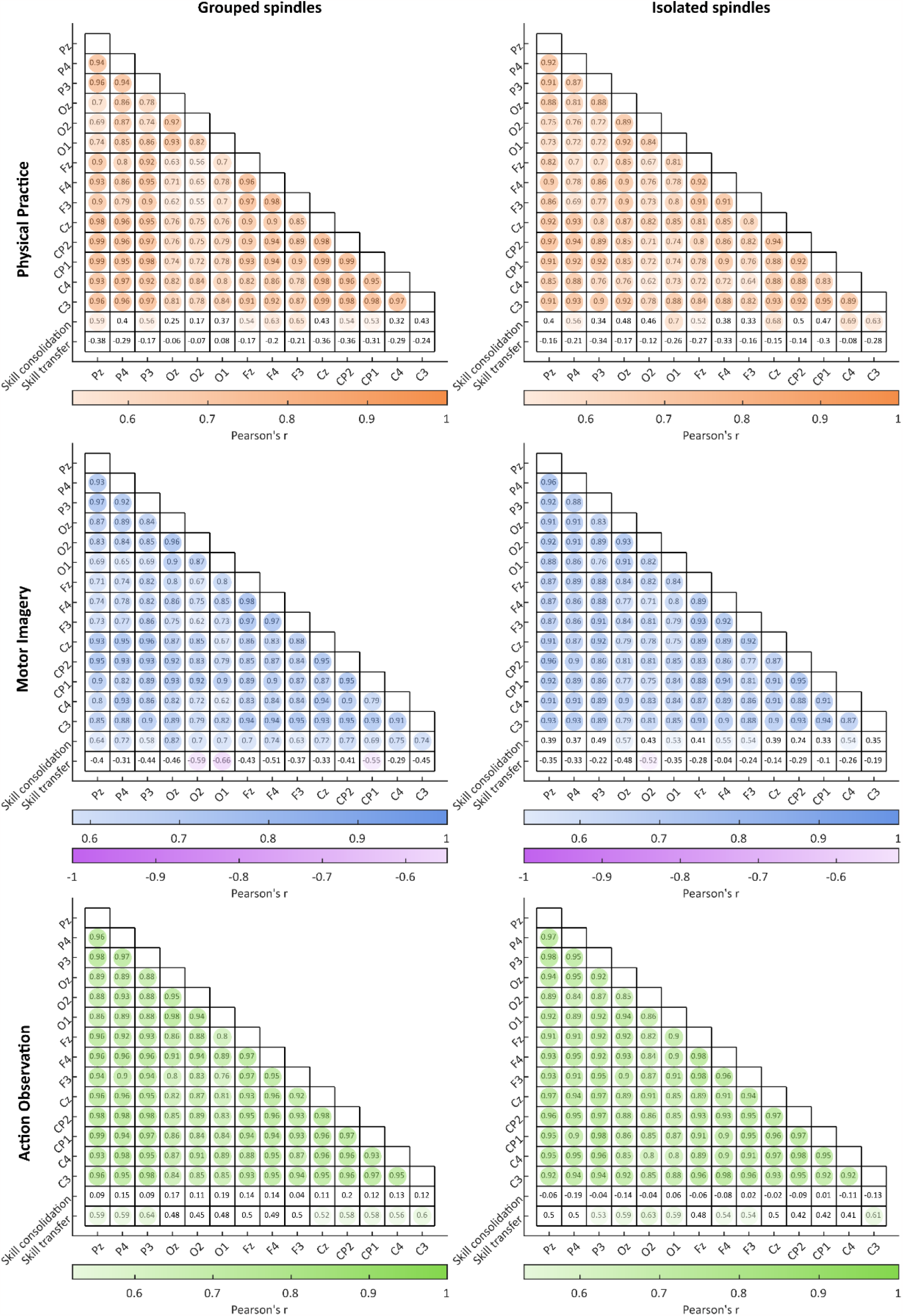
Grouped and isolated spindles in relation with skill consolidation and transfer. Relationship between the number of grouped spindles (left column) and isolated spindles (right column) detected over main scalp derivations with the magnitude of skill consolidation and transfer following physical practice (orange, first row), motor imagery (blue, second row) and action observation (green, third row). Pearson correlation coefficients (*r*) are reported for each correlation. Colored circles correspond to significant p-value (*p* < 0.05).

**Fig. S3.**
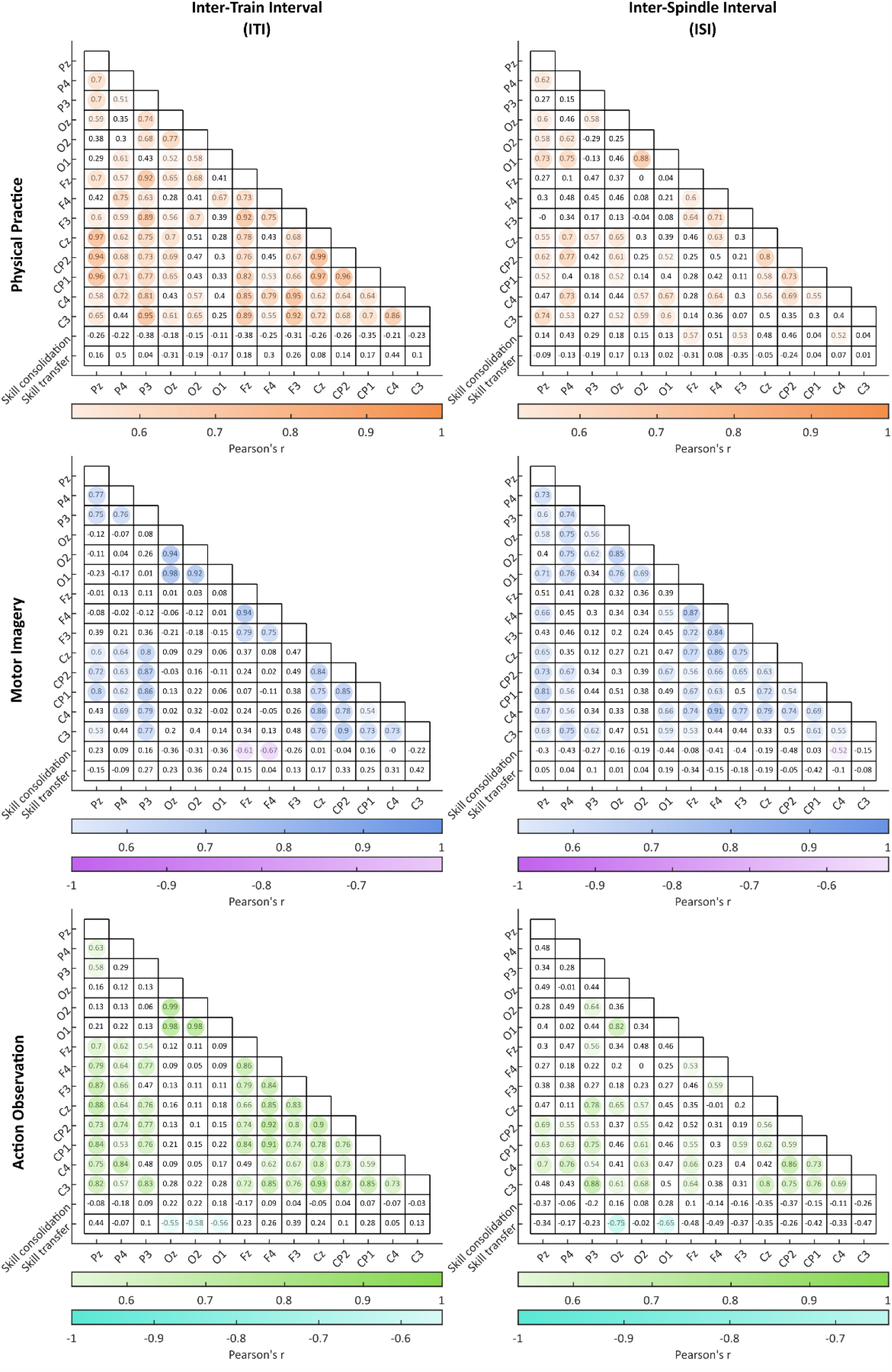
Sleep spindle clustering and rhythmicity in motor skill consolidation and transfer. Relationship between the inter-train interval (left column) and inter-spindle interval (right column) detected over main scalp derivations with the magnitude of skill consolidation and transfer following physical practice (orange, first row), motor imagery (blue, second row) and action observation (green, third row). Pearson correlation coefficients (*r*) are reported for each correlation. Colored circles correspond to significant p-value (*p* < 0.05).

**Note S4. Multiple regression analyses of grouped and isolated spindles regarding skill consolidation and transfer**. Multiple regression analyses were separately performed for each group to determine whether grouped or isolated sleep spindles predict motor skill consolidation, with respect to the modality of practice. The multiple regression model significantly predicted skill consolidation for the MI group only (F(2,12) = 4.24, *p =* 0.04, *R*² = 0.41). More specifically, only the factor grouped spindles was found to significantly contribute to the prediction of the model (t_Grp_ = 2.32, *p =* 0.039; t_Iso_ = -0.35, *p =* 0.74), thus revealing that the higher the number of grouped spindles during the nap following MI practice, the higher the magnitude of skill consolidation. The multiple regression model failed to significantly predict skill consolidation in the PP group (F(2,12) = 3.32, *p* = 0.07, *R*² = 0.36) and the AO group (F(2,12) = 0.32, *p =* 0.73, *R*² = 0.05). Additional multiple regression analyses were separately performed for each group to determine whether grouped or isolated sleep spindles predict motor skill transfer capacities, with respect to the modality of practice. The multiple regression model failed to predict skill transfer capacity in the PP (F(2,12) = 1.03, *p =* 0.39, *R*² = 0.15), MI (F(2,12) = 1.24, *p =* 0.32, *R*² = 0.17), and AO groups (F(2,12) = 3.21, *p =* 0.08, *R*² = 0.35).

